# Myosin1D is an evolutionarily conserved determinant of animal Left/Right asymmetry

**DOI:** 10.1101/267146

**Authors:** Thomas Juan, Charles Géminard, Jean-Baptiste Coutelis, Delphine Cerezo, Sophie Polès, Stéphane Noselli, Maximilian Fürthauer

## Abstract

The establishment of Left/Right (LR) asymmetry is fundamental to animal development. While the pathways governing antero-posterior and dorso-ventral patterning are well conserved among different phyla, divergent mechanisms have been implicated in the specification of LR asymmetry in vertebrates and invertebrates. A cilia-driven, directional fluid flow is important for symmetry breaking in numerous vertebrates, including zebrafish^1–10^. Alternatively, LR asymmetry can be established independently of motile cilia, notably through the intrinsic chirality of the acto-myosin cytoskeleton^11–18^. Here we show that MyosiniD (Myo1D), which has been previously identified as a key determinant of LR asymmetry in Drosophila^*12,13*^, is essential for the formation and the function of the zebrafish LR Organizer (LRO). We show that Myo1D controls the polarity of LRO cilia and interacts functionally with the Planar Cell Polarity (PCP) gene VanGogh-like2 (Vangl2)^19^, to promote the establishment of a functional LRO flow. Our findings identify Myo1D as the first evolutionarily conserved determinant of LR asymmetry, and show that functional interactions between Myo1D and PCP are central to the establishment of animal LR asymmetry.

---

Numerous studies have revealed an intriguing diversity in the strategies establishing laterality both within and across phyla, making the identification of a unifying mechanism elusive^14,20,21^. An attractive hypothesis however, is that the chirality of actin filaments may provide a conserved template for organismal LR asymmetry^14,18^. Accordingly, actin-binding proteins govern molecular^22^ and cellular chirality^23^ and actin-dependent processes regulate invertebrate and vertebrate laterality^12–14,17,24^. Studies in Drosophila have identified the actin-based molecular motor protein Myosin1D (Myo1D, a.k.a. Myosin31DF) as an essential determinant of LR asymmetry^12,13,25^. Here, we study the contribution of this conserved factor to vertebrate laterality, by analyzing its function in zebrafish.

We first determined whether zebrafish and Drosophila Myo1D have conserved activities by testing the ability of fish *myo1D* to rescue the laterality defects of fly *myo1D* mutants. The two orthologous proteins show 69% sequence similarity. Remarkably, expressing zebrafish *myo1D* in Drosophila fully restored the chirality of genitalia rotation (Fig. 1a), a prominent LR marker in Drosophila^12,13^. These data indicate that zebrafish Myo1D is capable of mediating a conserved LR symmetry-breaking activity.

**Figure 1:**
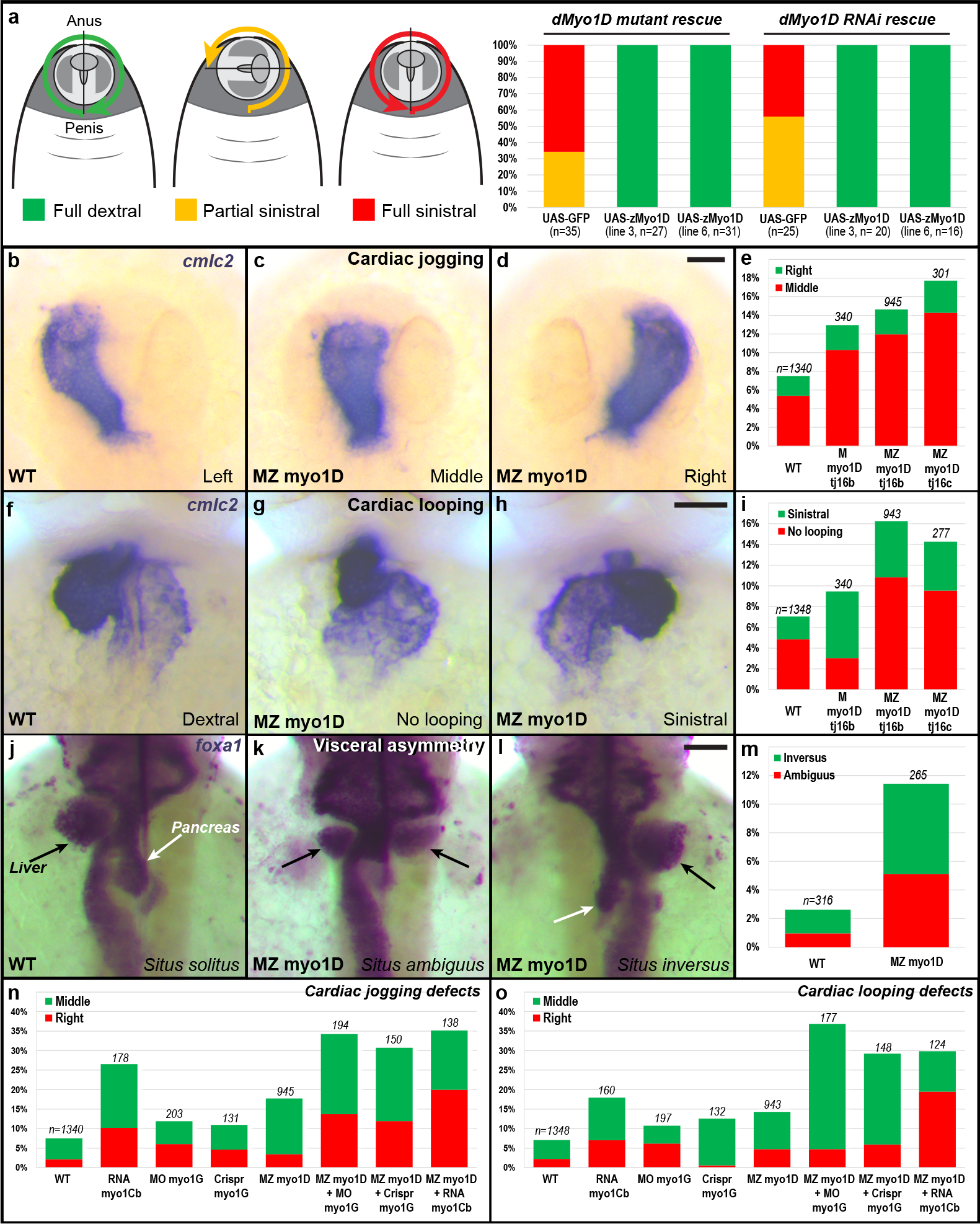
*myo1D* controls zebrafish Left/right asymmetry. **a,** Transgene-mediated expression of zebrafish *myo1D (zmyo1D)* restores the laterality of genitalia rotation in drosophila *myo1D (dmyo1D)* mutant or *dmyo1D* RNAi flies. Lines 3 and 6 are two independent transgenic insertions. **b-e,** Zebrafish MZ myo1D mutants present defects in leftward cardiac jogging. **b-d,** Dorsal views of the *cmlc2*-expressing heart primordium at 26 hpf, anterior up. **e,** Quantification of cardiac jogging defects in maternal (M) and maternal zygotic (MZ) *myo1D*^*tj16b*^ or *myo1D*^*tj16c*^ mutants. **f-i,** Cardiac looping is affected in MZ *myo1D* mutants. **f-h,** Frontal views of the *cmlc2*-expressing heart at 48 hpf, dorsal up. **i,** Quantification of cardiac looping defects in M *myo1D*^*tjl6b*^ and MZ *myo1D*^*tj16b*^ or *myo1D*^*tj16c*^ mutants. **j-m,** MZ *myo1D* mutants present defects in the leftward looping of the gut and the asymmetric development of the liver (black arrows) and pancreas (white arrows). **j-l,** Dorsal views of *foxal*-expressing visceral organs at 48 hpf, anterior up. **n,o,** Overexpression of zebrafish *myo1Cb* RNA or knock-down of *myo1G* by Morpholino (MO) or Crispr/Cas9 nuclease (Crispr) impair cardiac jogging (**n**) and looping (**o**). An increased frequency of defects is observed when these reagents are injected into MZ *myo1D* mutants. Scale bars: 30 μm.

To study zebrafish *myo1D* function, we generated frameshift mutations that disrupt the P-loop required for Myo1D motor activity^26^ and that delete the actin- and cargo-binding domains (Extended Data Fig. 1a). Homozygous mutants lacking zygotic *myo1D* function develop normally and give rise to fertile adults (Extended Data Fig. 1b). *In situ* hybridization reveals that, in addition to their zygotic expression, *myo1D* transcripts are maternally provided (Extended Data Fig. 1c-e). We therefore used homozygous mutant females to generate embryos lacking either only maternal (M) or both maternal and zygotic (MZ) *myo1D* activities. M and MZ *myo1D* mutants display LR asymmetry defects at the level of the heart (Fig. 1b-i), viscera (Fig. 1j-m) and brain (Extended Data Fig. 2a-f). Identical phenotypes were observed using two different mutant alleles (Fig. 1e,i). Similar defects are moreover generated when a morpholino that blocks the translation of both maternally deposited and zygotically transcribed *myo1D* RNA is injected in wild-type embryos (Extended Data Fig. 2g-m). Altogether, our experiments identify Myo1D as a conserved regulator of laterality in both vertebrates and invertebrates.

In Drosophila, Myosin1C (Myo1C, a.k.a. Myo61F) acts as an antagonist of the essential dextral determinant Myo1D^27^. *In flies, myo1C* overexpression phenocopies *myo1D* loss-of-function. However, Drosophila *myo1C* is itself dispensable for LR asymmetry^27^. The zebrafish genome encodes two *myo1C* homologues, *myosinlCa and myosinlCb (myo1Ca & b)*. Like in Drosophila, zygotic *myo1Ca* mutants and MZ *myo1Cb* mutants do not display LR defects (not shown). Interestingly, Myo1Cb overexpression using mRNA microinjection causes cardiac laterality defects (Fig. 1n,o), thus mimicking *myo1D* loss-of-function, as observed in Drosophila^27^.

We noticed that the overexpression of zebrafish *myo1Cb* enhances MZ *myo1D* mutant LR defects (Fig. 1n-o), raising the question whether additional Myo1D-like activities might contribute to the regulation of LR asymmetry. Genome analysis shows that the zebrafish Myosin1G (Myo1G) protein is closely related to Myo1D (79% similar amino acids). While *myo1G* knock-down through morpholino or Crispr/Cas9 nuclease injection elicits modest cardiac laterality defects (Fig. 1n,o), the injection of these reagents into MZ *myo1D* mutant embryos allows to increase the frequency of LR defects to a level equivalent to the one observed upon *myo1Cb* overexpression (Fig. 1n,o). Altogether, these findings provide evidence that Myo1D-like agonists (Myo1D & G) and their antagonist Myo1Cb form an evolutionarily conserved pathway for the control of LR asymmetry.

The observation that MZ *myo1D* mutants present laterality defects at the level of the heart, viscera and brain (Fig. 1b-m, Extended Data Fig. 2a-f), suggests that *myo1D* may control the formation and/or function of the central zebrafish LR Organizer (LRO), Kupffer’s Vesicle (KV)^3,28^. The ciliary LRO flow induces the expression of both the Nodal ligand *southpaw (spaw)* and the transcription factor *pitx2* in the left lateral plate mesodem^29,30^. Consistent with a potential role of Myo1D at the level of the LRO, the asymmetric expression of these two genes is disrupted in MZ *myo1D* mutants (Fig. 2a-f). To further test Myo1D’s function in the LRO, we looked at the asymmetric expression of the TGFβ antagonist *charon/dand5*, which is the earliest known transcriptional response to the KV flow^4,31^. While weak symmetric *dand5* expression is initiated prior to the establishment of a functional KV, *dand5* asymmetry arises as the LRO matures, peaks by the 8-somite stage and decreases again during later development (Fig. 2g-m). In accordance with a dysfunction of the LRO, *dand5* asymmetry is reduced in MZ *myo1D* mutants and completely lost in MZ *myo1D* mutants injected with *myo1Cb* RNA or *myo1G* morpholino (Fig. 2n-t). These data demonstrate that prior to the first morphological manifestations of organ LR asymmetry, Myo1D is required at early stages for the LRO-dependent establishment of asymmetric gene expression.

**Figure 2:**
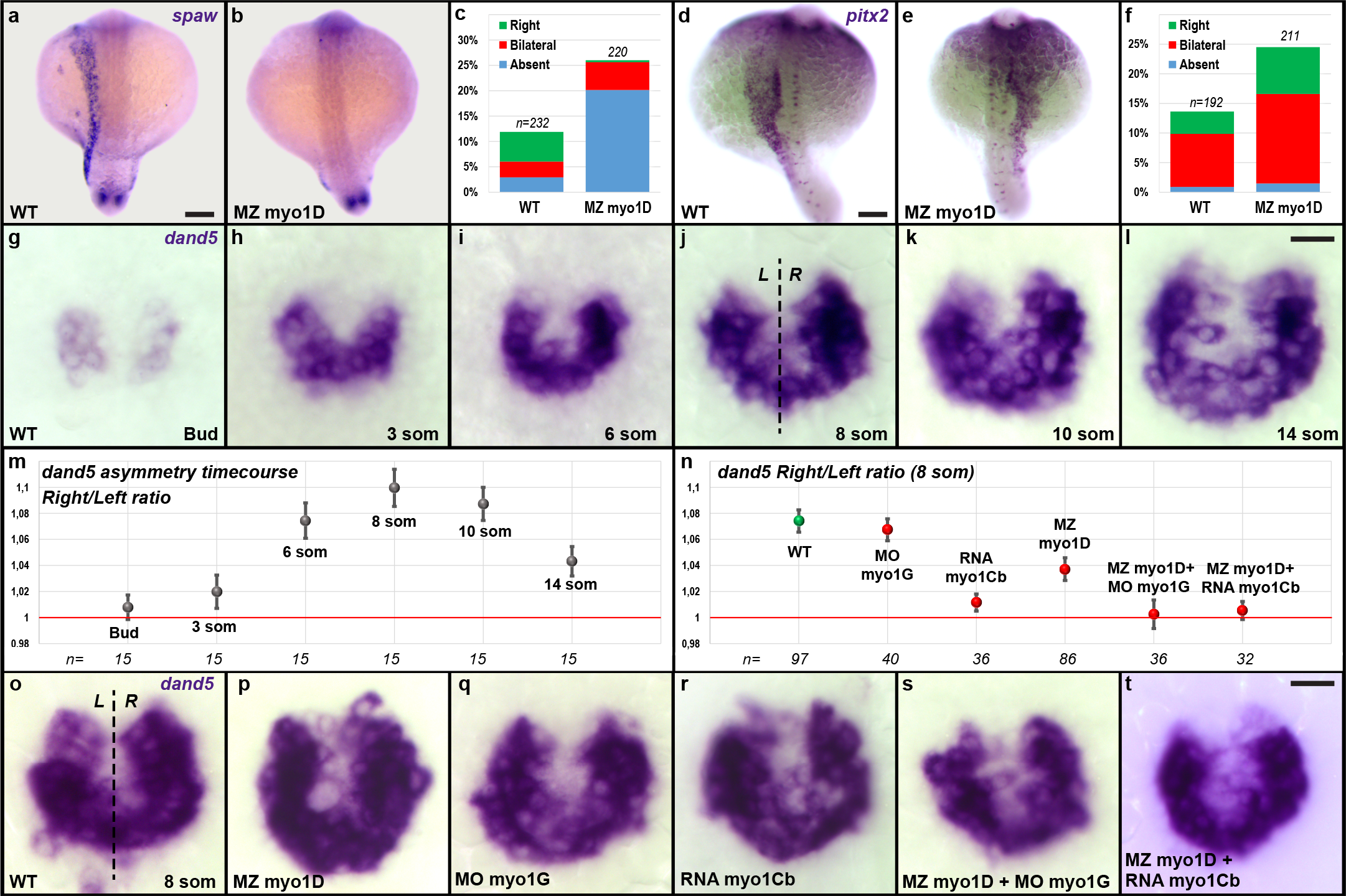
*myo1D* is required for Left/Right Organizer function. **a-f,** MZ *myo7D* mutants display an impaired expression of *southpaw (spaw)* and *pitx2* in the left lateral plate mesoderm. **a,b,d,e,** Dorsal views of 18-somite stage embryos, anterior up. **c,f,** Quantifications of *spaw* (**c**) and *pitx2* (**f**) expression. **g-m,** Temporal evolution of the asymmetry of *dand5* expression between the end of gastrulation (Bud) and the 14-somite stage. **m,** Quantifications of the relative abundance of *dand5* transcripts on the Left (L) and Right (R) side of WT ABTÜ embryos reveal a peak of LR asymmetry at the 8-somite stage. **n-t,** The 8-somite stage asymmetry of *dand5* asymmetry is reduced in MZ *myo7D* mutants and completely lost upon additional morpholino knock-down of *myo1G* (MO myo1G) or overexpression of *myo1Cb* (RNA myo1Cb). **g-l** and **o-t** are dorsal views of the LRO, anterior up. Error bars in **m** and **n** indicate SEM. Scale bars: 50 μm in **a,b,d,e**. 20 μm in **g-l** and **o-t**.

To further dissect the role of *myo1D* in zebrafish LR asymmetry, we analyzed its contribution to the formation and function of the KV. While *myo1D* is dispensable for the migration and coalescence of sox17-positive KV precursor cells (Extended Data Fig. 3), we found that the average KV size and cilia number are reduced in MZ *myo1D* mutants (Fig. 3a,b,e,g-i). As LRO function requires a minimal organ size and number of cilia^4,32^, we monitored KV size, cilia number and laterality in individual embryos. No correlation could be observed between organ size, cilia number and the appearance of laterality defects (Fig. 3e, Extended Data Fig. 4a). Accordingly, treatment with IBMX and Forskolin, which promote KV lumen inflation^33^, rescues KV size but not laterality defects in MZ *myo1D* mutants (Fig. 3c-f). While the loss of *myo1D* causes a modest reduction of cilia size (Fig. 3g,h,j), it has no incidence on their motility or their clustering in the anterior KV half, two factors known to be required for LRO flow (Extended Data Fig. 4b,c)^34^.

**Figure 3:**
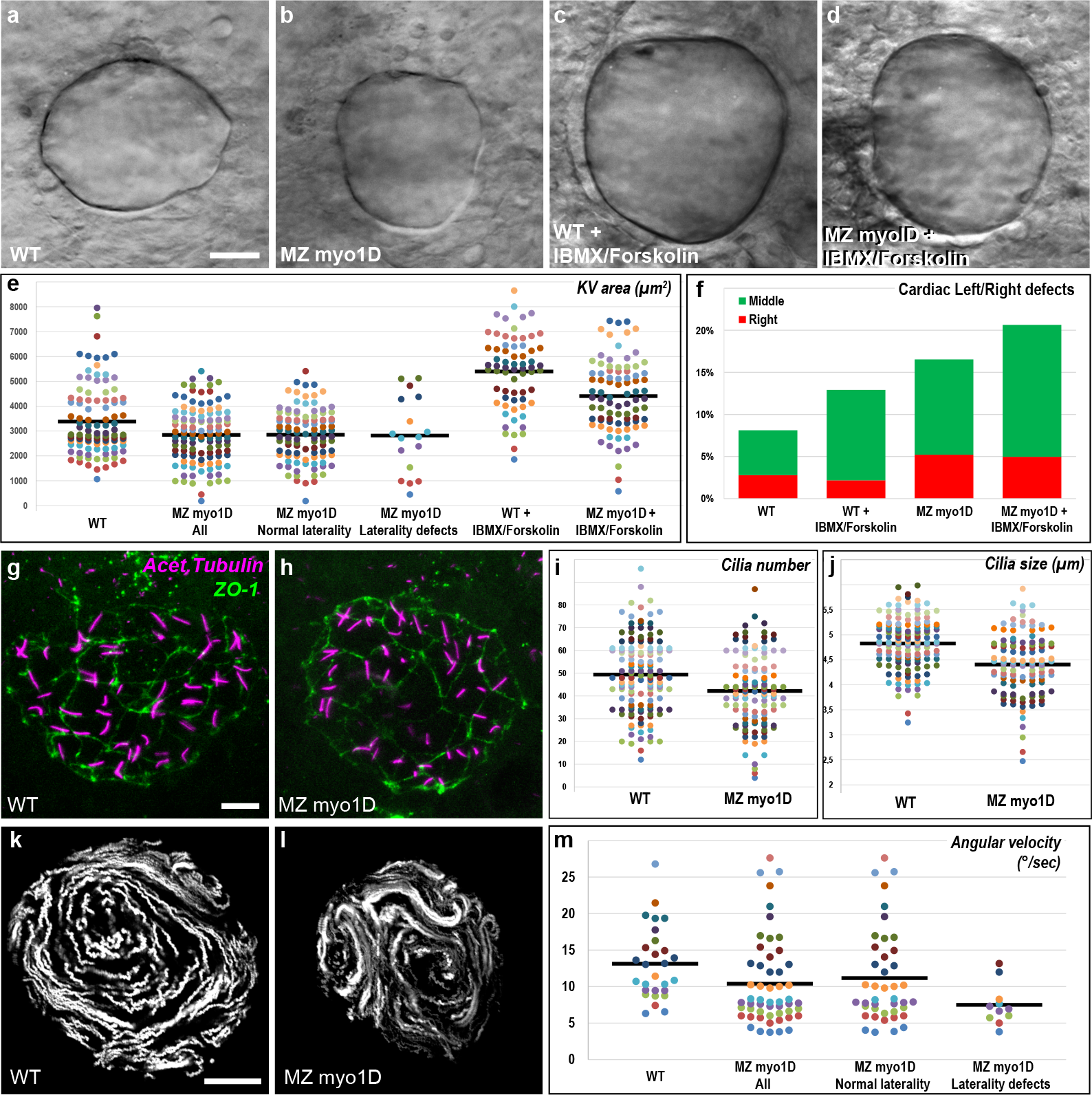
The establishment of the ciliary Left/Right Organizer flow requires *myo1D*. **a-e,** Compared to WT (n=104), MZ *myo1D* mutants (n=118) present a reduced KV size. Treatment with IBMX and Forskolin promotes KV lumen inflation and increases organ size in WT (n=66) and MZ *myo1D* mutants (n=85). **e,** Dot plots of KV equatorial surface area in individual embryos. Note that MZ *myo1D* mutant laterality defects occur independently of KV size. **f,** IBMX/Forskolin treatment restores KV size (**d,e**), but not laterality in MZ *myo1D* mutants. **g-j,** Cilia number and size are reduced in MZ *myo1D* mutants. **g,h,** Projection of images from confocal stacks used to quantify number and length of cilia (acety-lated tubulin, magenta) in the KV (ZO-1 positive cells, green) in WT (**g**) and MZ *myo1D* mutants (**h**). **i,j,** Dot plots representing the number and average length of cilia in individual embryos (WT n=141 embryos/6967 cilia; mutant n=117/4939). **k-m,** KV flow is altered in MZ *myo1D* mutants. **k,l,** Temporal projections of the trajectories of fluorescent microspheres in WT (n=28) and MZ *myo1D* mutants (n=53). **m,** Dot plots of mean angular flow velocity in individual embryos. Note that laterality defects correlate with low flow velocites. **a-d,g,h,k,l** are dorsal views of 8-somite stage KVs, anterior up. Horizontal grey bars in **e,i,j,m** represent mean values of different data sets. Scale bars: 20 μm in **a-d,k,l**. 10 μm in **g-h**.

To visualize the LRO flow, we tracked fluorescent microspheres in the KV lumen. While wild-type controls mostly displayed circular trajectories (Fig. 3k), MZ *myo1D* mutants often presented aberrant flow patterns (Fig. 3l). To quantitatively assess the LRO flow, we determined its mean angular velocity as a measure of the effective circular flow strength (see Methods). Results show that MZ *myo1D* mutants display a decrease in angular flow velocity (Fig. 3m). Importantly, LR asymmetry defects in MZ *myo1D* mutants correlate with pronounced reductions in flow velocity (Fig. 3m). In contrast, treatment of control wild-type embryos with Ouabain to pharmacologically reduce KV size leads to an increase in flow velocity with no associated LR defects (Extended Data Fig. 5). These data show that Myo1D is required for the establishment of a functional LRO flow, independently of KV size.

How does Myo1D control KV flow? It is known that embryos with no, or immotile, cilia lack a detectable flow^3,4^. In MZ *myo1D* mutants, cilia are motile but the LRO flow is erratic, a phenotype reminiscent of mutants for the Planar Cell Polarity (PCP) pathway component *vangl2* (Fig. 4a-c)^19 35^. We confirmed this phenotypic similarity through quantitative analysis of KV flow in *vangl2*^*m209*^ mutants (Fig. 4e,f). Previous work showed that the establishment of LR asymmetry requires the generation of peak flow velocities in the anterior KV^4,34^. We found that both MZ *myo1D* and *vangl2* mutants display a reduction in overall angular velocity (Fig. 4e) and a disruption of the antero-posterior flow pattern (Fig. 4f). Interestingly, while *myo1D* and *vangl2* mutants have similar flow phenotypes, removal of one copy of *vangl2* reduces the penetrance of LR defects in M or MZ *myo1D* embryos (Fig. 4g), suggesting that the two genes affect the LRO in opposite ways.

**Figure 4:**
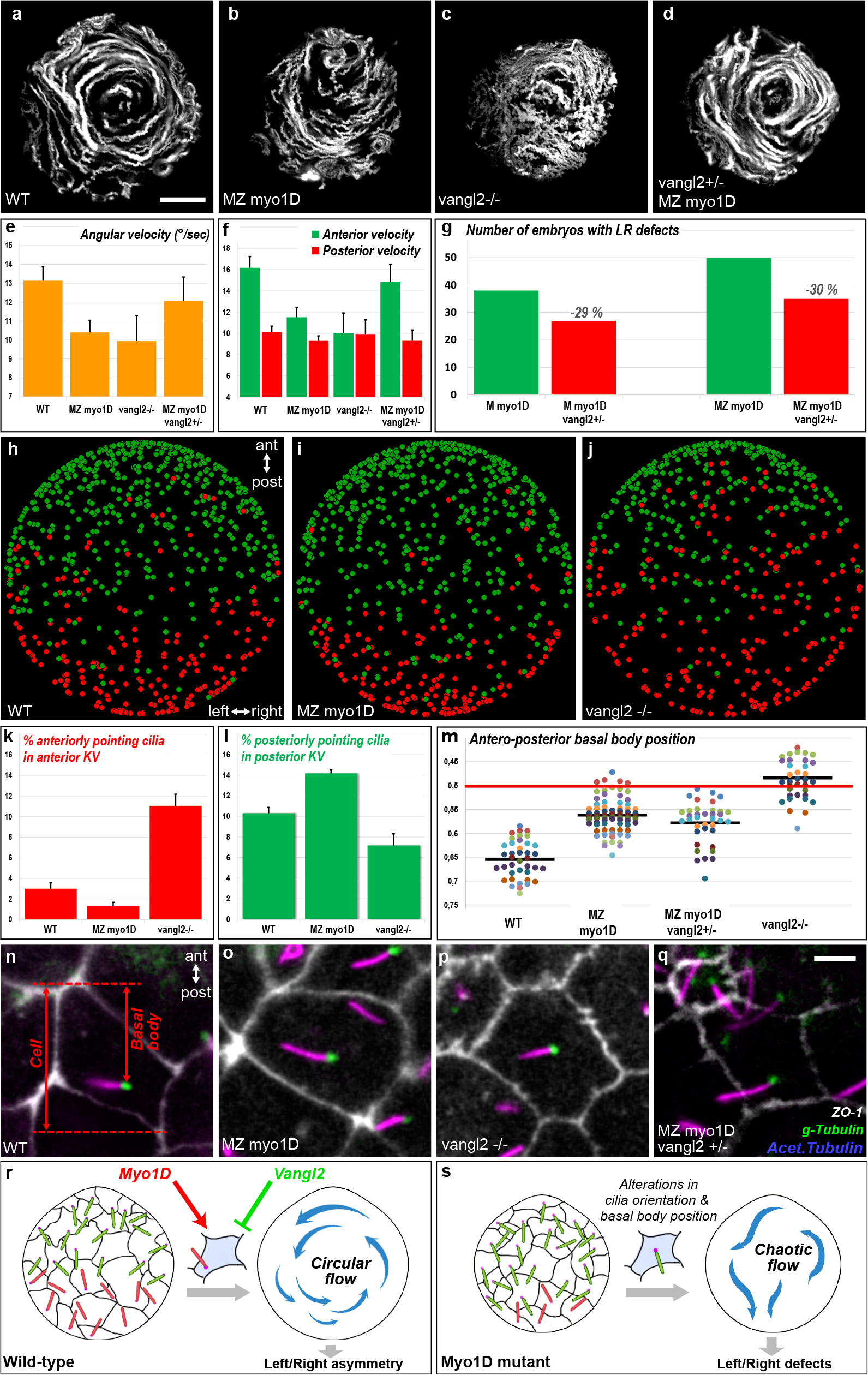
*myo1D* interacts with *vangl2* to control the morphogenesis and function of the Left/Right Organizer. **a-d,** Temporal projections of trajectories of fluorescent microspheres in the KV lumen. Compared to WT (n=28), MZ *myo1D* (n=53) and *vangl2* (n=21) mutants present a similar impairment of LRO flow. Flow pattern is partially restored in MZ *myo1D;vangl2*+/− (n=20). **e,f,** MZ *myo1D* and *vangl2* mutants display an impairment in flow velocity (**e**) and antero-posterior pattern (**f**), both of which are partially restored in MZ *myo1D;vangl2+/−*. **g,** The removal of one copy of *vangl2* reduces the occurrence of LR defects in maternal (M) or maternal zygotic (MZ) *myo1D* mutants (see Methods). **h-l,** MZ *myo1D* and *vangl2* mutants display opposite defects in KV cilia orientation. **h-j** represent the position of cilia pointing to the anterior (red) or posterior (green) in WT, MZ *myo1D* and *vangl2* mutants (see Extended Data Fig.6 and Methods). *vangl2* mutants present an excess of anteriorly pointing cilia (**k**), while more posteriorly pointing cilia are present in MZ *myo1D* (**l**). **m-q,** *myo1D* and *vangl2* ares required for basal body positioning. **m,** Dot plot representing the average antero-posterior basal body position in individual embryos (see Methods). Compared to WT (n=40 embryos/744 basal bodies), basal bodies are displaced anteriorly in MZ *myo1D* (n=74/1521) and *vangl2* (n=36/526) mutants, but partially restored to their posterior position in MZ *myo1D;vangl*+/− (n=37/826). Grey bars represent mean values. **n-q,** High magnification views of KV cells (ZO-1) with cilia (acetylated tubulin) and basal bodies (γ-Tubulin). All data collected at 8-somite stage. Anterior is up in **a-d,h-j,n-q**. Error bars in **e,f,k,l** indicate SEM. Scale bars: 20 μm in **a-d**. 5 μm in **n-q**. **r,s,** Schematic representation of the role of Myo1D in the KV.

PCP signaling controls the posterior positioning of the ciliary basal body and the orientation of the ciliary rotation cone to promote LRO flow^19,36–38^. Recently, rat *myo1D* has been shown to control cilia orientation in multiciliated epithelia^39^. We therefore imaged cilia in live embryos and found that zebrafish *myo1D* and *vangl2* both affect cilia orientation: While *vangl2* mutants present an excess of anteriorly pointing cilia in the anterior KV, the loss of *myo1D* increases the number of posteriorly pointing cilia in the posterior organ half (Fig.4h-l, Extended Data Fig. 6). Additionally, the inactivation of either *myo1D* or *vangl2* causes an anterior displacement of the basal body (Fig. 4m-p). The positioning of the basal body is however partially restored when one copy of *vangl2* is removed in MZ *myo1D* mutants (Fig. 4m,q). These findings suggest that *myo1D* and *vangl2* affect PCP-dependent KV cyto-architecture, but do so in opposite ways, consistent with our genetic interaction data (Fig. 4g). We therefore reasoned that if the observed cilia defects are relevant for LRO function, then a reduction of *vangl2* gene dosage should restore KV flow in MZ *myo1D* mutants. Accordingly, the removal of one copy of *vangl2* allows to partially rescue KV flow pattern and velocity in MZ *myo1D* mutants (Fig. 4d,e,f).

Taken together, our data show that Myo1D is essential for the formation and function of the zebrafish LRO, identifying the first common determinant of LR asymmetry in both vertebrates and invertebrates. Our work also reveals that a functional interaction between zebrafish Myo1D and PCP controls the positioning and orientation of KV cilia that are required for the establishment of a directional, symmetry-breaking flow (Fig. 4r,s). Interestingly, recent work has shown a functional link between Myo1D and PCP pathways in Drosophila^40^ and in Xenopus (M. Blum, personal communication). We propose that the coupling between Myo1D and PCP polarity systems represent an evolutionarily conserved genetic framework controlling LR asymmetry in animals.

## ACKNOWLEDGEMENTS

The present project was supported by the ANR project grants DRO-ASY (ANR-13-BSV2-006, S.N.) and DroZeMyo (M.F., S.N.). The work in the lab of M.F. was supported by a CNRS/INSERM ATIP/Avenir 2010 grant and an HFSP Career Development Award (00036/2010). T.J. benefited from a PhD fellowship from INSERM/Région PACA/Cancéropôle PACA and the Labex SIGNALIFE (ANR-11-LABX-0028-01). We thank S.Lopes, B.Ciruna and T.Lepage for the sharing of reagents and R.Rebillard for excellent fish care. Confocal microscopy and timelapse imaging of the KV fluid flow were performed at the iBV PRISM imaging platform with the help of Magali Mondin.

## AUTHOR CONTRIBUTIONS

The genetic analysis of MyosinI function in zebrafish LR asymmetry was entirely performed by T.J. J.B.C. performed initial morpholino knock-down experiments of zMyo1D and zMyo1G. S.P. cloned zMyo1D and G, analyzed Myo1D expression and performed Myo1D morpholino knock-down and Myo1Cb overexpression experiments together with T.J. D.C. generated zMyo1D-transgenic Drosophila lines. C.G. performed the rescue experiments of the fly Myo1D mutants with zMyo1D. S.N. and M.F. designed the study, analyzed the data and wrote the manuscript.

## AUTHOR INFORMATION

The authors declare no competing financial interests. Correspondence and request for materials should be addressed to S.N. or M.F.

## METHODS

### Fly strains and genetics

Flies were grown according to standard procedures. For inter-specific rescue experiments, the *UAS::zMyo1D* transgene (Zebrafish Myo1D ORF cloned into pUAST vector) was expressed either in *myo1D* depleted flies (*w*^*1118*^ *; ptc::GAL4, myo1D*^*k1*^ *; UAS::dmyo1D-RNAi* ^13^) or in a *myo1D*^*k2* 13^ null mutant background (*da::GAL4*; *myo1D*^*k2*^)(*da::GAL4*, Bloomington Drosophila Stock Center #55851). Two independent insertions of the *UAS::zMyo1D* transgene (line 3 and 6) were used and showed similar results.

Rotation of the male genitalia was scored as previously described^42^. Briefly, male genital plates are scored according to the angle made between the dorso-ventral and anus-penis axes when viewed from the posterior. Full dextral/clockwise (+360°) corresponding to the wild-type situation; partial sinistral/counterclockwise rotation (from −1 to −359°); full sinistral/counterclockwise rotation (−360°).

### Zebrafish strains and embryo maintenance

Embryo were raised in 0.3x Danieau medium (17.4 mM NaCl, 0.21 mM KCl, 0.12 mM MgSO_4_, 0.18 mM Ca(NO3)_2_, 1.5 mM Hepes, pH 7.6) at 28,5°C, and staged according to standard criteria^43^. If necessary, 1-phenyl-2-thiourea (Sigma) was added at 30 mg/l to prevent embryonic pigmentation.

### Crispr/Cas9 Mutagenesis

Crispr/Cas9 mutagenesis of zebrafish *myo1D* (Ensembl gene ENSDARG00000036863) was performed in a wild-type TÜ background. gRNA design and in vitro transcription of Cas9 RNA were performed according to reported protocols^44^. gRNA was transcribed from a template oligo as previously described^45^. The sequence of the selected Crisp target site was 5’-GAGTGGAGCTGGAAAAACAG**AGG**-3’ (bold lettering indicates the PAM motif). The efficiency of Crispr/Cas9-induced mutagenesis was monitored at 24 hpf using a T7 endonuclease assay^44^ on a PCR amplicon comprising the Crispr target region (Forward primer: 5’-TCTTCACTGACACTGGTATG-3’, Reverse primer: 5’-CCATCACTGCAGCAGAAATGAGAG-3’). Adult F0 fish were outcrossed to AB wild-type fish and DNA extracted from F1 progeny. Mutations were identified through direct sequencing of the same PCR amplicon that was used in the T7 endonuclease assay.

Zebrafish *myo1G* (Ensembl gene ENSDARG00000036104) was inactivated through Crispr/Cas9 mutagenesis at the target site 5’-GATGTCATTGAGGACTACAG**GGG**-3’. A pre-assembled complex of purified Cas9 protein (NEB, 954 ng/μl) and *myo1G* gRNA (200 ng/μl) was injected into one cell stage embryos and the cutting efficiency monitored using a T7 endonuclease assay on a *myo1G* PCR amplicon (Forward primer: 5′-GGAGAAGTAGTGGTGTCCGTTAAC-3’, Reverse primer: 5’-CTCACTTTTGGGCTAACAGCTC-3’).

### Fish strains and molecular genotyping

The following lines were used for this work: Tg(−5.0*sox17*:EGFP)zf99^41^, Tg(*cmlc2*:RFP)^46^, *vangl2*^m209 47^.

Wild-type and mutant alleles of *myo1D* were identified through allele-specific PCRs. The allele-specific forward primers 5’-TGGAGCTGGAAAAAGGCTCGT-3’ (*myo1D*^**tj16b**^) and 5’-GTGGAGCTGGAAAAAGGCTATAC-3’ (*myo1D*^*tj16c*^) were used together with the generic reverse primer 5’-CCATCACTGCAGCAGAAATGAGAG-3’ to genotype for the presence of different *myo1D* mutant alleles. PCR amplicon size was 133 bp for *myo1D*^*tj16b*^ and 145 bp for *myo1D*^*tj16c*^. The wild-type *myo1D* allele was amplified using the forward primer 5’-AGAGTGGAGCTGGAAAAACAGA-3’ and the reverse primer 5’-CCCATCCCTCGTGTGAAACTAAATCAC-3’ to yield a 339 bp amplicon. PCR amplification with GoTagG2 polymerase (Promega) was performed using the following cycling conditions: Initial denaturation 2min 95°C; 10 cycles [30s 95°C, 30s 65°C first to 55°C last cycle, 30s 72°C]; 20 cycles [30s 95°C, 30s 55°C, 30s 72°C]; final extension 5min 72°C.

To generate maternal zygotic *myo1D* mutants (MZ *myo1D*^*tj16b/tj16b*^ or MZ *myo1D*^*tj16c/tj16c*^), *myo1D* heterozygous fish were incrossed and the progeny raised to adulthood. PCR genotyping of these animals allowed identifying homozygous mutant as well as homozygous wild-type fish which were then used to produce MZ *myo1D*^*tj16b/t16b*^ or MZ *myo1D*^*tj16c/tj16c*^ mutant embryos, as well as wild-type *myo1D*^*+/+*^ control embryos. *myo1D*^*tj16b*^ or *myo1D*^*tj16c*^ homozygous mutant animals presented similar laterality defects (Fig. 1e,i). Unless mentioned otherwise, experiments were performed using the *myo1D*^*tj16b*^ allele.

For the genetic interaction experiment displayed in Fig. 4g, *myo1D*^*tj16b*^ homozygous mutant females were crossed with *myo1D*^*tj16b/+*^; *vangl2*^*m209/+*^ males. A total of 912 embryos derived from 6 independent crosses were scored and molecular genotyping performed for all embryos presenting cardiac laterality defects. Genotyping for the *vangl2*^*m209*^ mutant allele was performed as previously described^48^.

### Plasmid generation

The *myo1Cb* ORF was amplified from a mixed stage pool of cDNAs using the primers: 5’-**GATCCCATCGATTCGA**CGGAATCCGGGTCATGATG-3’ and 5’-**AGGCTCGAGAGGCCTT**GGTCACGACTCGTCCATC-3’ and cloned into the pCS2+ vector. Bold letterings indicate primer overhangs that are also present in the pCS2+ sequence and were used to recombine insert and vector using Gibson assembly (NEB). A similar strategy was used to clone the *myo1D* ORF into pCS2+ using the primers 5’-**GATCCCATCGATTCGA**CAGCTTATTATGGCAGAACACG-3’ and 5’-**AGGCTCGAGAGGCCTT**AAATGGGATCCTGGTCCTCTAG-3’.

### RNA and morpholino injections

*myo1Cb*-pCS2+ was linearized with KpnI, *myo1Cb* RNA *in vitro* synthesized using the mMessage mMachine SP6 transcription kit (Ambion) and the resulting RNA injected at 50ng/μl. RNA encoding Arl13b-GFP was made as previously described^19^ and injected at 20ng/μl. Two translations-blocking morpholinos (Gene Tools) were used: Myo1D, 5’-ACTTTCGTGTTCTGCCATAATAAGC-3’ (injected at 750 μM); Myo1G, 5’-CCTCCAGCTCCGCCATCCCTGCAAC-3’ (62.5 μM). All reagents were injected at the one cell stage at a volume of 0.88nl together with 0.2% phenol red.

### RNA in situ hybridizations

Whole mount RNA in situ hybridizations were performed as previously described^49^. The following probes have been previously described: *cmlc2*^50^, *foxa1/fkd7*^51^, lefty1^52^, *pitx2*^30^, *sox17*^53^, *southpaw*^29^, *dand5/charon*^31^. To generate a *myo1D* probe, *myo1D* was amplified from *myo1D*-pCS2+ using the following primers 5’-CGC**TAATACGACTCACTATAGGGAGA**GTTCTAGAGGCTCGAGAGG-3’ and 5’-GGCAAGACCAAAGTTTTCATCCGC-3’. The *in situ* probe was then synthesized directly from the PCR product using the incorporated T7 promoter (bold lettering). *In situ* hybridizations were documented using a Leica M205FA stereomicroscope equipped with an Infinity2-5C color CCD (Lumenera).

### Immunohistochemistry

Dechorionated embryos were fixed overnight at 4°C in PEM (80 mM Sodium-Pipes, 5 mM EGTA, 1 mM MgCl_2_) − 4% PFA − 0.2% TritonX100. After washing 2 × 5 min in PEMT (PEM - 0.2% TritonX100), 10 min in PEM - 50 mM NH4Cl, 2 × 5 min in PEMT and blocking in PEMT - 2%BSA, embryos were incubated 2 hrs at room temperature with primary antibodies. Following incubation, embryos were washed during 5, 10, 15 and 20 min in PEMT, blocked in PEMT - 2%BSA, and incubated again with secondary antibodies for 2 hrs. Embryos were again washed during 5, 10, 15 and 20 min in PEMT, mounted in PEM-0,75% LMP-Agarose and analyzed on a Zeiss LSM710 laser scanning confocal microscope. The following primary antibodies were used in this study: Mouse@ZO-1 (1:250, Invitrogen 33-9100), Mouse@Acetylated Tubulin (1:2000, Sigma T7451) and Rabbit@γTubulin (1:250, Sigma T5192). The secondary antibodies Goat@MouseIgG1-Alexa568, Goat@MouseIgG2b-Alexa647 and Goat@Rabbit-Alexa488 were all obtained from Invitrogen and used at a dilution of 1:500.

### Pharmacological treatments

Stock solutions of IBMX (100mM, Sigma) and Forskolin (10mM, Sigma) were prepared in DMSO. Ouabain (1mM, Sigma) was prepared in distilled water. Embryos were treated with 10 μM Forskolin and 40 μM IBMX, or 5 μM Ouabain at bud stage and washed in Danieau medium after imaging at the 8-somite stage.

### Microinjection and tracking of fluorescent beads in the Left/Right Organizer

Fluoresbrite Polychromatic Red Microspheres 0.5μm (Polysciences) were manually injected into the KV lumen. The movement of fluorescent beads was imaged on a wide field Zeiss Axio Observer Z1 microscope with a 40X EC Plan-Neofluar PH/DCII NA 0.75 dry objective using an Andor Neo sCMOS camera and a DsRed filter. In this setup we obtained images with a pixel width of 0.1625 μm and recorded the KV flow for 2 minutes with a frame interval of 0.04s. Automated tracking of fluorescent beads was performed using a custom-made ImageJ script (available on request). The macro generates an automated tracking file that is compatible with the ImageJ MTrackJ plugin. Following automated detection, tracks were manually curated using MTrackJ to correct for any errors of the tracking algorithm and the tracking parameters exported in a .CSV datasheet for further analysis. Using this procedure, an average of 41624 tracking points were obtained per embryo. KV images were oriented precisely along the antero-posterior axis based on the position of the notochord as well as the characteristic indentation that is found at the anterior pole of the KV in the region were cells are most closely packed^54^. The center of the KV was determined as the point lying midway between the anterior and the posterior extremity of the KV on a line that extends posteriorly from the anterior pole of the KV. The angular flow velocities displayed in Figures 3 and 4 were calculated using the center of the KV as the point of origin of a polar coordinate system.

### Quantification of ciliary orientation

KV motile cilia were labeled using Arl13b-GFP RNA microinjection and imaged on a Zeiss710 laser scanning confocal microscope. The entire KV was scanned with z-sections of 0,2 μm distance and a time interval of 0.48s between two consecutive planes. Movies were then treated in ImageJ with a Mean 3D filter (x radius=1, y radius=1, z radius=10) to visualize the ciliary rotation cone and be able to orient it from base to tip (center of the cone). Each cilium was drawn on the KV using the ImageJ arrow tool. Cilia positions and angles were then extracted from the arrow ROIs using a custom-made ImageJ script. To compare the position and orientation of cilia in different embryos as shown in Fig. 4h-j, the relative positions of different cilia were normalized in the common space of a circular virtual KV. Each cilium was placed on the virtual KV according to its relative radial position with respect to the organ border. ImageJ scripts are available on request.

### Quantification of the antero-posterior positioning of the ciliary basal body

Confocal Z-stacks of the entire KV were acquired with a distance of 0,6 μm between two consecutive planes. The KV was oriented along the antero-posterior axis based on the position of the notochord. Basal bodies were identified as γ-Tubulin positive spots that are located at the base of an Acetylated Tubulin positive ciliary axoneme. The junctional marker ZO-1 was used to outline KV cells and determine the antero-posterior extent of each cell. We restrained our analysis to cells for which the entire outline could be unambiguously identified in the ZO-1 channel. As the KV is an ellipsoid structure, this applies essentially to the cells of the dorsal roof and ventral floor of the KV. Based on this criterium, we were able to determine the antero-posterior position of an average of 19.4 basal bodies per embryo. The anteroposterior location of the basal body was quantified by calculating the ratio between the distance of the basal body from the anterior-most limit of the ZO-1 signal and the full antero-posterior length of a cell inferred from the ZO-1 signal. According to this convention, a value of 0 would indicate a basal body that is located at the anterior extremity of the cell, while a value of 1 would correspond to an extreme posterior location.

### dand5/charon quantification

*dand5/charon in situ* hybridization pictures were analyzed with ImageJ. A bounding rectangle was fit on the *dand5* expression domain and the mean grey levels of the left and right part of the rectangle measured. Dand5 Right/Left enrichment was determined as the ratio between right and left mean grey levels as previously described^55^.

---

**Extended Data Figure 1:**
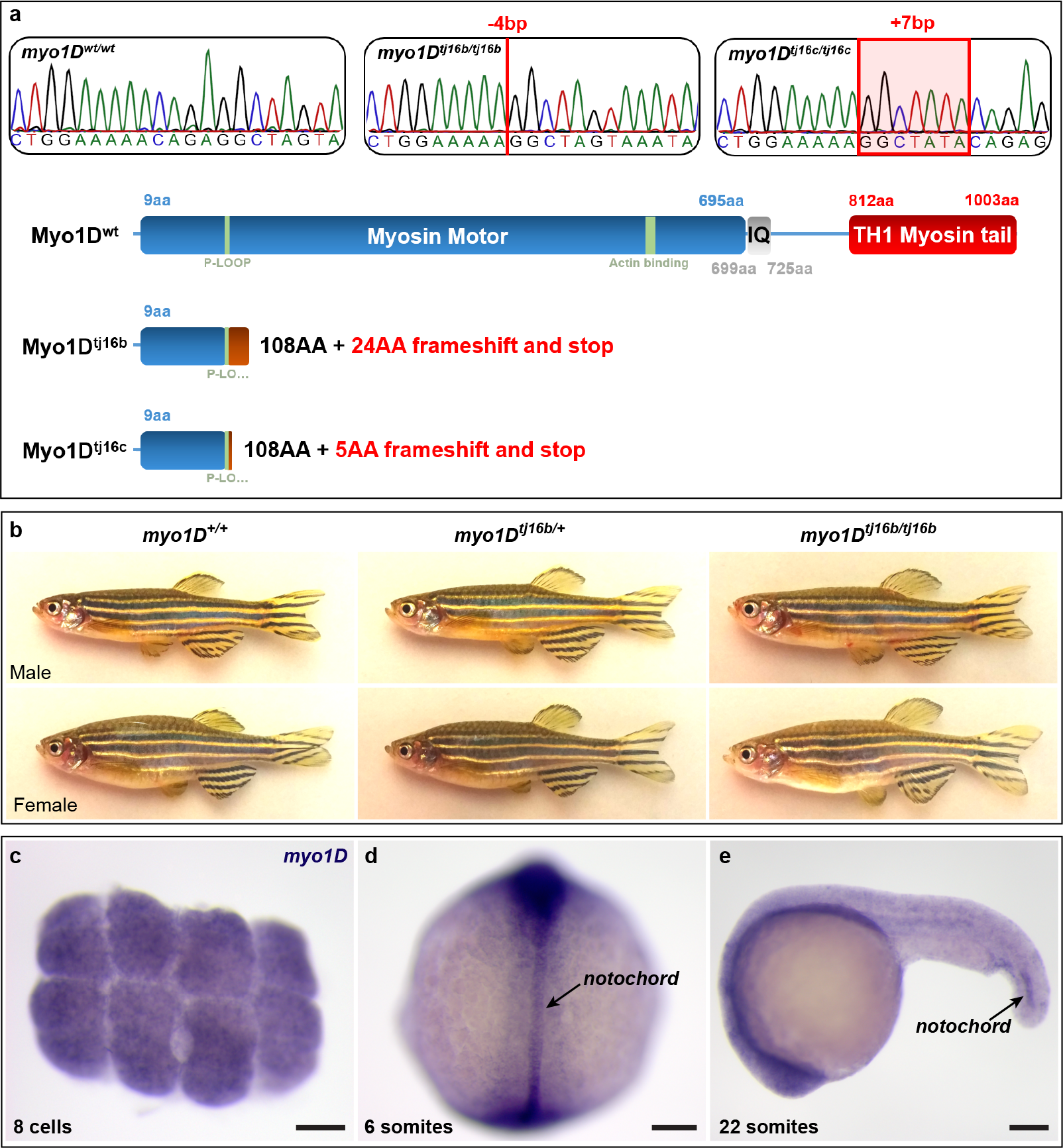
Genetic inactivation of zebrafish *myo1D*. **a,** Sequence chromatograms and schematic representation of *myo1D* mutants generated by Crispr/Cas9 mutagenesis. **b,** *myo1D* homozygous mutant adult fish are indistinguishable from heterozygous or homozygous wild-type siblings. **c,** Whole-mount *in situ* hybridization reveals that *myo1D* transcripts are maternally supplied and can be detected at the 8-cell stage (animal pole view), well before the activation of the zygotic genome. **d, e,** Zygotic *myo1D* transcripts are present at low levels throughout the embryo. At segmentation stages, increased transcript levels are detected in the notochord. **d,** dorsal view, anterior up. **e,** lateral view, anterior to the left. Scale bars: 100 μm.

**Extended Data Figure 2:**
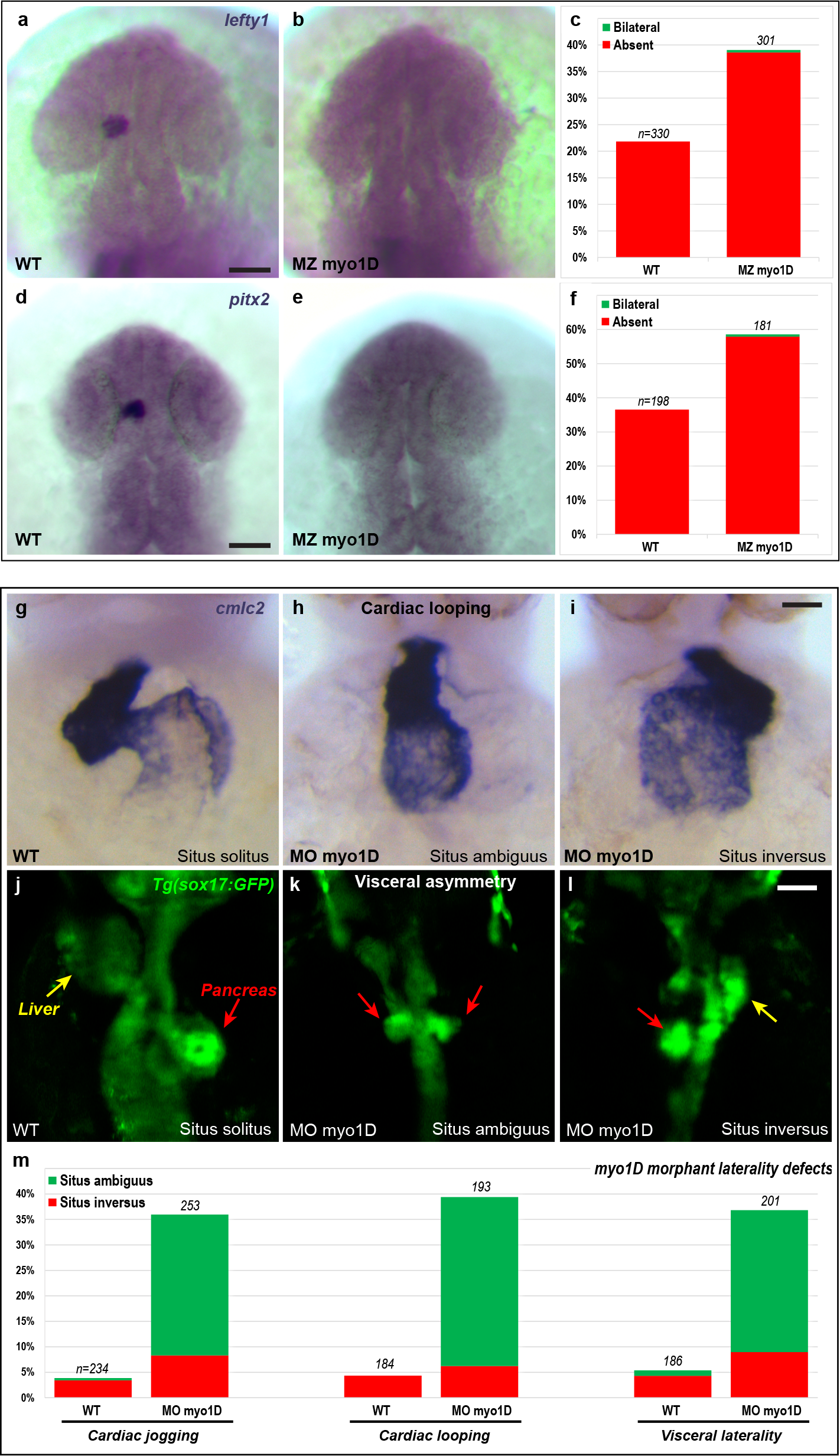
Zebrafish *myo1D* controls Left/Right asymmetry in the brain, heart and viscera. **a-f,** Asymmetric expression of *lefty1* and *pitx2* is impaired in the brain of MZ *myo1D* mutant animals. Dorsal views of *lefty1* (22-somite stage) and *pitx2* (25-somite stage) expression in WT (**a,d**) and MZ *myo1D* mutant (**b,e**) embryos. Quantification of *lefty1* (**c**) and *pitx2* (**f**) expression shows that asymmetric gene expression often fails to be established in mutants. **g-m,** Morpholino knock-down of *myo1D* elicits laterality defects at the level of the heart and viscera. *myo1D* morphants present defects in cardiac jogging (**m**) and looping (**g-i,m**). **g-i** are frontal views of the **cmlc2**-expressing heart at 48 hpf, dorsal up. **j-m,** *myo1D* knock-down impairs the leftward looping of the gut as well as the lateralized development of the liver (yellow arrows) and pancreas (red arrows). Dorsal views of visceral organs highlighted by an endodermal *sox17:GFP* transgene^41^. Anterior is up. Scale bars: **a,b,d,e** 30 μm. **g-i** 20 μm. **j-l** 50 μm.

**Extended Data Figure 3:**
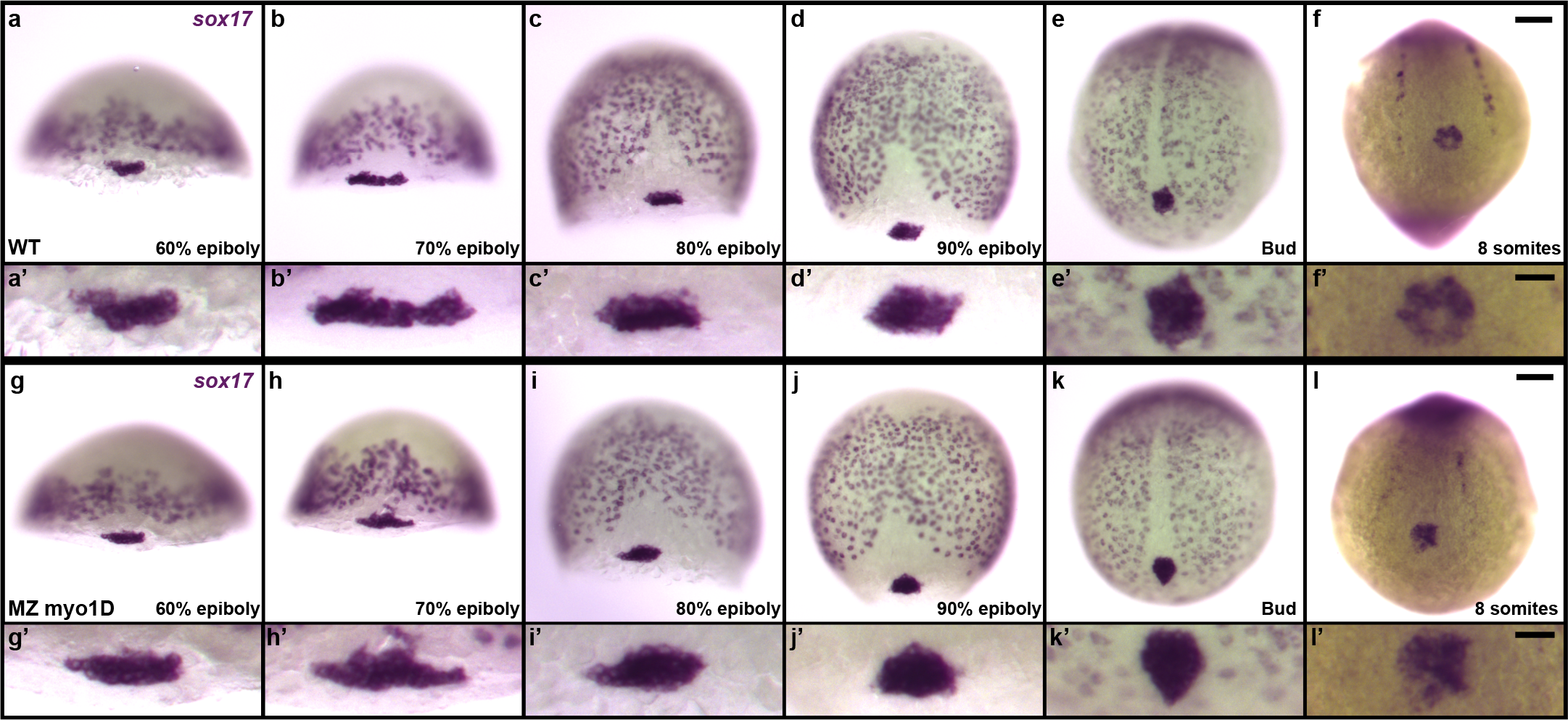
Zebrafish *myo1D* is not required for the specification and migration of Left/Right organizer precursor cells. **a-l,** *sox17 in situ* hybridization marks scattered endodermal cells as well as the posterior cluster of dorsal forerunner cells that give rise to the fish Left/Right Organizer, Kupffer’s Vesivle (KV). Comparison of WT (**a-f**) and MZ *myo1D* mutant (**g-l**) embryos reveals no differences in the number and behavior of KV precursor cells. **a-d,g-j,** are dorsal views, anterior up. **e,f,k,l,** are vegetal views of the tail bud region at the end of gastrulation (**e,k**) and the 8-somite stage (**f,l**). Anterior is up. The upper pictures represent low magnification views of the whole embryo. The lower panels show high magnification views of the KV precursor cells. Scale bars: **a-l**, 50 μm. **a’-l’,** 20 μm.

**Extended Data Figure 4:**
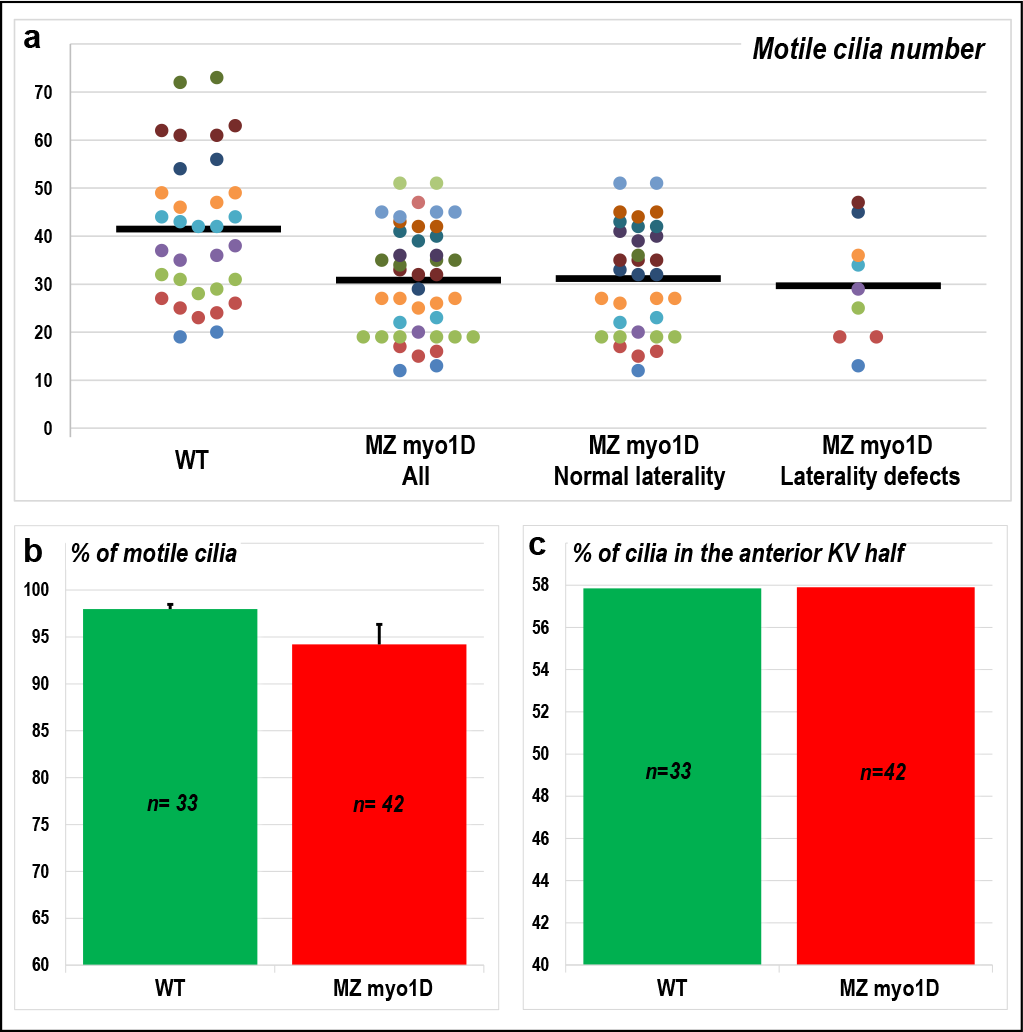
Zebrafish *myo1D* is dispensable for ciliary motility. **a-c,** High speed confocal imaging of *Arl13b-GFP* labelled cilia was used to analyze the number (**a**), motility (**b**) and positioning (**c**) of KV cilia. **a,** Dot plot representing the number of motile cilia in individual embryos. MZ *myo1D* mutants display a lower cilia number, but this decrease does not correlate with the occurrence of laterality defects. Horizontal grey bars indicate the mean values for different data sets. **b,** *myo1D* loss of function does not impair ciliary motility. Error bars indicate SEM. **c,** In both WT and MZ *myo1D* mutant animals, the majority of cilia are located in the anterior half of the KV. All data were collected at the 8-somite stage.

**Extended Data Figure 5:**
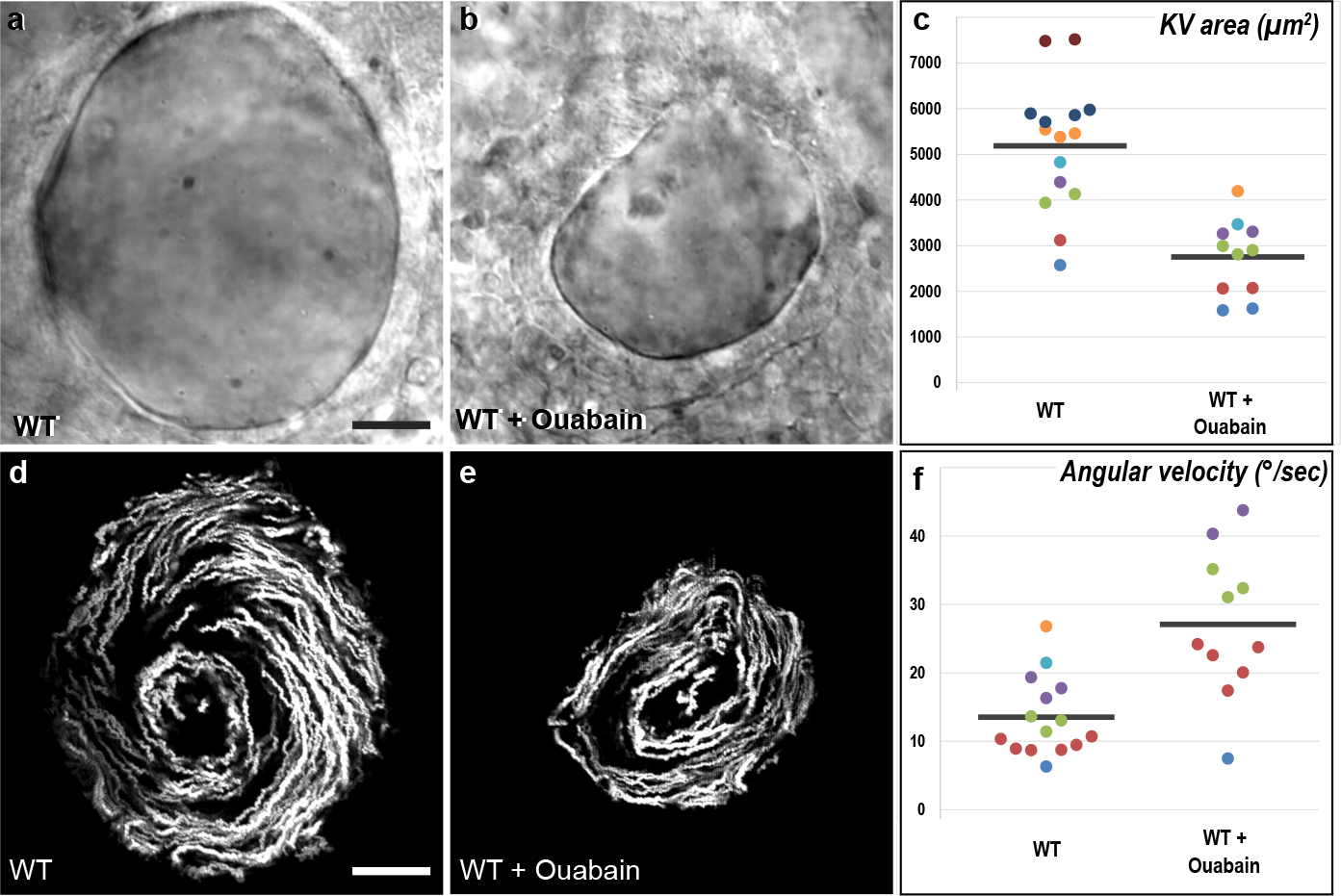
The velocity of the zebrafish Left/Right Organizer flow increases upon pharmacological reduction of organ size. **a-c,** Ouabain treatment of WT embryos inhibits lumen inflation of Kupffer’s Vesicle (KV) and thereby reduces organ size. **a,b,** are brightfield images of the equatorial plane of control (**a,** n=15) and Ouabain-treated embryos (**b,** n=11). **d-f,** Both WT control (**d**) and Ouabain-treated embryos (**e**) display a circular flow pattern. **d,e** are temporal projections of the trajectories of fluorescent microspheres in the KV lumen of the embryos displayed in **a** and **b. f,** The angular velocity of the KV flow is increased upon Ouabain treatment. **a,b,d,e** are dorsal views of 8-somite stage KVs, anterior is up. Horizontal grey bars in **c,f** represent mean values for different data sets. The WT control embryos displayed in this figure are also part of the data set that is used in **Fig. 3m**. Scale bars: 20 μm.

**Extended Data Figure 6:**
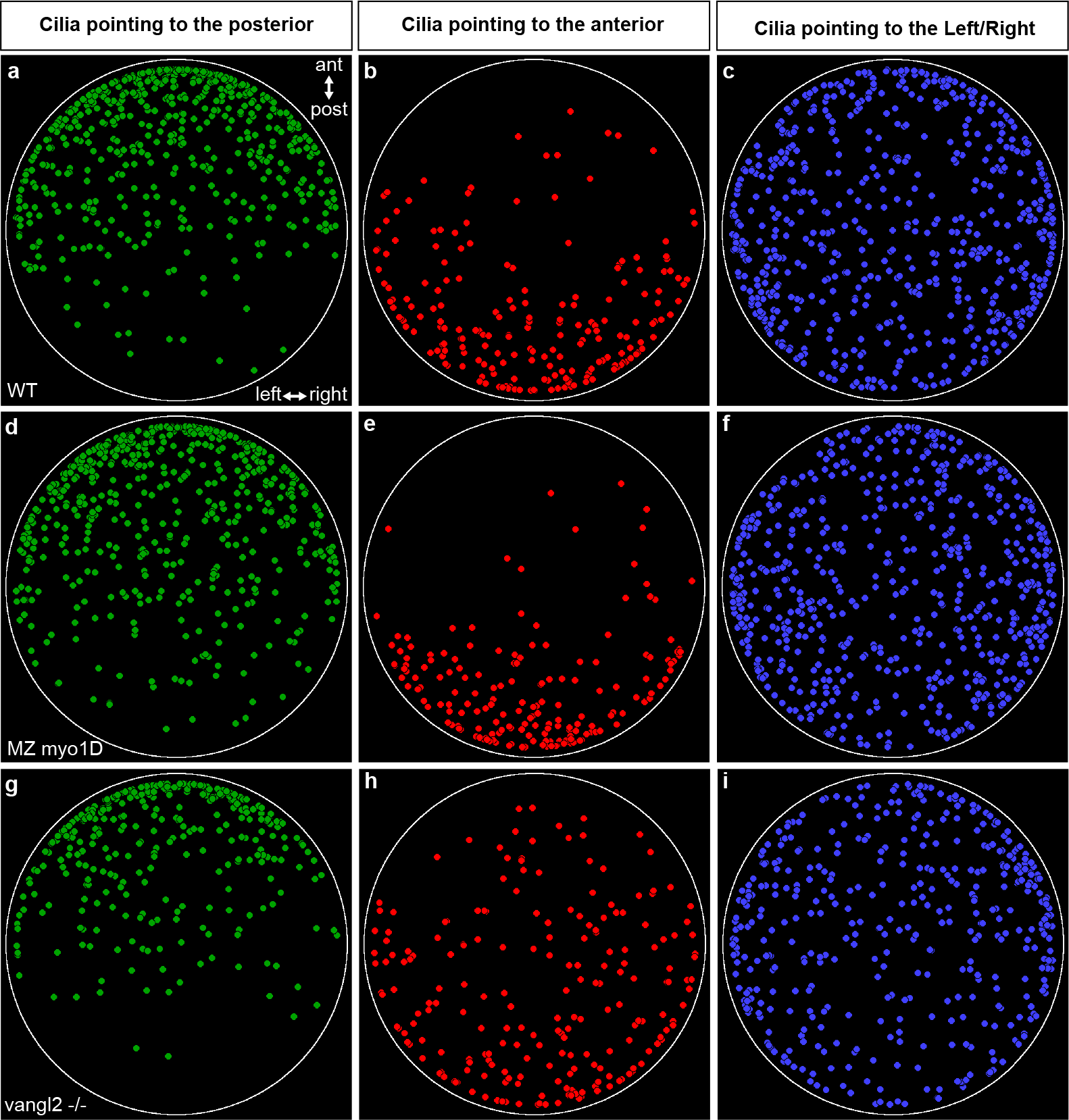
Zebrafish *myo1D* and *vangl2* have opposite effects on cilia orientation in the Left/Right Organizer. **a-i,** The orientation of ciliary rotation cones in Kupffer’s Vesicle (KV) was determined using high speed confocal imaging of *Arl13b-GFP* labelled embryos (see Methods). 2-dimensional dot plots indicate the position of cilia within the KV of WT (n=33 embryos/1369 cilia), MZ *myo1D* (n=42/1296) and *vangl2* (n=31/966) mutant embryos. Cilia were categorized according to their orientation: Green dots indicate cilia that point towards the posterior side of the KV with a deviation from the antero-posterior axis of no more than 45°. Red dots indicate cilia that point towards the anterior side of the KV with a deviation of no more than 45° from the antero-posterior axis. The remaining cilia (i.e. those pointing to the left or right organ wall) are indicated by blue dots. Each plot is a cumulative representation all the cilia of a given genotype in the space of a normalized virtual KV. Note that posteriorly pointing cilia invade the posterior half of the KV in MZ *myo1D* mutants (**d**), while an excess of anteriorly pointing cilia is present in the anterior KV half of *vangl2* mutants (**h**). The embryos in this figure correspond to the data set that is also displayed in **Fig. 4h-l**.

